# Isomeric lipid signatures reveal compartmentalised fatty acid metabolism in cancer

**DOI:** 10.1101/2021.11.01.466716

**Authors:** Reuben S. E. Young, Andrew P. Bowman, Kaylyn D. Tousignant, Berwyck L. J. Poad, Jennifer H. Gunter, Lisa K. Philp, Colleen C. Nelson, Shane R. Ellis, Ron M. A. Heeren, Martin C. Sadowski, Stephen J. Blanksby

## Abstract

1.0

Cellular energy and biomass demands of cancer drive a complex dynamic between uptake of extracellular fatty acids (FA) and *de novo* synthesis. Given that oxidation of *de novo* synthesised FAs for energy would result in net-energy loss, there is an implication that FAs from these two sources must have distinct metabolic fates - however hitherto FAs were considered part of a common pool. To probe FA metabolic partitioning, cancer cells were supplemented with stable-isotope labelled FAs. Structural analysis of the resulting glycerophospholipids revealed that *labelled* FAs from uptake were largely incorporated to canonical (*sn*-)positions on the glycerol backbone. Surprisingly, *labelled* FA uptake disrupted canonical isomer patterns of the *unlabelled* lipidome and induced repartitioning of *n*-3 and *n*-6 polyunsaturated-FAs into glycerophospholipid classes. These structural changes evidence differences in the metabolic fate of FAs derived from *uptake* or *de novo* sources and demonstrate unique signalling and remodelling behaviours usually hidden to conventional lipidomics.

**Highlights:**

- Lipid isomers reveal discrete metabolic compartmentalisation in cancer
- FAs derived from uptake and *de novo* synthesis have different metabolic fates
- Stearate uptake signals for PUFA (*n*-3 and *n*-6) repartitioning between lipid classes
- *sn*-positional isomers are a marker for aberrant lipid metabolism

## 2.0 Introduction

Although seemingly banal, the structure-function relationship of glycerophospholipids (GPL) is far more complex and insightful than first meets the eye. While the amphoteric nature of the molecule importantly allows for chemical partitioning, the individual functional groups serve numerous other specific purposes. The charged hydrophilic head group creates the possibility for lipids to be directed to particular subcellular compartments whereby the lipophilic FAs can impart variable physicochemical properties to the molecule and initiate signalling.^1^ Arguably, two of the most important processes in dictating the modelling and remodelling of mammalian GPL structure are the Kennedy pathway and the Lands cycle. While the Kennedy pathway describes the *de novo* synthesis of phosphatidylcholine (PC) and phosphatidylethanolamine (PE) from cytidine-diphosphate diacylglycerol,^2^ the Lands cycle chronicles the process by which phospholipase and acyltransferase enzymes remodel GPL-FAs.^3^ While these processes logically have implications in cellular energy production, whether *de novo* generated FAs or extracellular FAs (from uptake) have identical or varied roles is not well understood. Given that cancer cells are known to modulate and switch between *de novo* and uptake FA metabolism to opportunistically source energy for growth,^4-5^ understanding if subfractions of FAs serve discrete purposes is of importance when elucidating the molecular mechanisms of cancer. For example, a cancer cell actively increasing the production of FAs only to oxidise these to meet energy demands is counterproductive and thus the cell might instead be compelled to source extracellular FAs. For this to be possible however, there must be compartmentalisation and discrete metabolic treatment of *de novo* or uptake FAs – a sentiment that is contrary to the current idea that the cell has a common “pool” of FAs, used to meet metabolic demand.^4, 6^

One of the more important enzymes within the Lands cycle is phospholipase-A2 (PLA2), a lipase which is responsible for the enzymatic cleavage of the FA located at the *sn*-2 position within GPLs. Notably, PLA2 has been shown to exhibit a high affinity of action towards GPLs containing polyunsaturated FAs (PUFA), such as arachidonic acid (AA; 20:4*n*-6) and docosahexaenoic acid (DHA; 22:6*n*-3).^7^ Amongst other functions the release of AA and DHA in turn initiate pro- or anti-inflammatory responses through the cyclooxygenase and lipoxygenase pathways.^8-9^ Concurrent with the release of the *sn-*2 FA is the formation of a 1-acyl-2-lyso-GPL, or 2-lyso-GPL, with a hydroxyl group at the *sn*-2 position of the glycerol-backbone. While these lyso-species are themselves signalling mediators,^10^ they also readily undergo re-esterification, reverting to diacyl-GPLs via acyltransferase enzymes, such as the lyso-phosphatidic acid acyltransferase (LPAAT) or lyso-phosphatidylcholine acyltransferase (LPCAT) isoforms.^3, 11-12^ The labile nature of lyso-GPLs also allows physicochemical properties such as pH and temperature to influence a behaviour known as acyl chain migration, whereby a FA is able to migrate from one *sn-*position to a free hydroxyl group on the glycerol-backbone. Unlike the *sn-*selectivity of PLA2, acyltransferase enzymes are not reported to exhibit preference for 1-lyso or 2-lyso species and as such acyl chain migration that occurs prior to esterification will influence lipid molecular structure. Because Kennedy and Lands metabolic pathways affect different structural motifs of the GPL molecule, they are able to dictate both the transport of lipids around the cell and the initiation of signalling cascades through the release of biologically active FAs. This allows for independent organelle membrane modifications as well as the initiation of signalling cascades through the release of biologically active FAs. Although the various mechanisms and outcomes of lipid remodelling have been studied extensively, little is known about the impact that *sn-*isomeric lipid species have on remodelling and transport, or conversely, the impact these mechanisms have on lipid isomer populations. While it is now known that the GPLs of both prokaryotic and eukaryotic cells exhibit a preference for unsaturated FAs to be at the *sn*-2 position, prior to 2003 there was little-to-no indication that *sn-*positional isomers were present in eukaryotic cells.^13-14^ Ekroos *et al*. displayed that not only was there a quantifiable population of *sn-*1 unsaturated GPLs, but the relative abundance of these regioisomer populations can vary between mammalian GPL samples.^13^ This was exemplified in more recent studies into mammalian GPL regioisomers, which revealed that bovine liver,^15^ egg yolk and sheep kidney^16^ exhibited a stronger preference for canonical *sn*-positions than synthetically prepared standards.

Although an independent lipase enzyme, phospholipase A1 (PLA1), is known to catalyse the cleavage of FAs in the *sn-*1 position of specific GPL classes,^17^ the preference for unsaturated FAs to exist in the *sn*-2 position is thought to be strongly linked with Lands cycle lipid remodelling and PLA2 functionality.^7, 9, 18^ This enhanced *sn-*positional specificity in biological systems implies that cellular intervention is required for maintenance. Studies into the peroxidation of unilamellar liposomes comprised of specific phosphatidylcholine *sn-*isomers (containing palmitic acid and linoleic acid) revealed oxidation rates were *sn*-isomer dependent.^19^ Using radiolabelled FAs in mouse models, others showed that while the supplemented cis-FA (oleic acid) showed esterification specificity to the *sn*-2 position of GPLs, the trans-FA species (elaidic acid) was equally esterified to *sn*-1 and -2 positions. More recently, molecular dynamics simulations revealed that unsaturated FAs at the *sn*-1 position of GPLs create more ordered (and hence less fluid) membranes than their unsaturated *sn*-2 counterparts.^20^ These functional differences may help explain variations in *sn-*isomer distribution that are observed to exist in recent tissue models. In a pivotal study, Paine *et al*. showed that lipid isomers were tightly regulated and highly organised within the grey and white matter structures of brain tissue, and were subsequently disrupted in murine cancer tumours.^21^ Therefore, because lipid species containing saturated FAs at the *sn*-1 and unsaturated FAs at the *sn-*2 are the typical structures identified as the majority isomer in lipid canon, to aid with explanation throughout, these isomers will be referred to as ‘canonomers’. Conversely, lipids displaying atypical or apocryphal *sn-*isomeric structure (with the unsaturated FA contained at the *sn*-1 position) will be referred to ‘apocromers’.

Using stable isotope labelled FAs and a prostate cancer cell line (LNCaP), here we trace the metabolic fate of isotopically labelled FAs in cellular lipids, specifically tracking the *sn*-position (*sn-*position) at which they are incorporated into glycerophospholipids (GPL). Using a tandem mass spectrometric technique that combines collision-induced dissociation (CID) and ozone-induce dissociation (OzID; *i*.*e*., CID/OzID), we determine that lipids incorporating FAs from supplementation have isomer profiles that are distinct from lipids incorporating *de novo* synthesised (or previously stored) FAs. Monitoring the distribution of *sn-*isomers across resected LNCaP xenograft tumours (from mice) using isomer-resolved molecular imaging reveals that apocromeric species are spatially correlated to PUFA containing GPLs and lyso-lipids. Treating mice with a potent and selective phospholipase A2 Group IIA (sPLA2-IIA) inhibitor (KH064), and conducting the same correlative analysis reveals that spatial correlation between apocromer, PUFA-GPLs and lyso-lipids is weakened. Given that there are multiple known changes to the physico-chemical environment of cancer tumours^22-23^ and lyso-lipid acyl chain migration is known to be affected by pH and temperature,^24^ we propose that lipid apocromer species are the product of lipid remodelling in a perturbed physico-chemical environment and show potential as a biomarker for aberrant lipid metabolism in cancer tumours.

## 3.0 Results

### 3.1 FA origin and its influence on isomeric lipid remodelling

To investigate the impact of FA supplementation, LNCaP cells were either nil supplemented (NS) or supplemented with ^12^C16-16:0 (palmitic acid; 12PA), ^13^C16-16:0 (palmitic acid; 13PA) or ^13^C18-18:0 (stearic acid; 13SA). Using conventional lipidomics methods, the GPL sum composition for each supplement was determined, giving total GPL class abundance (Fig. 1A-B). Given the mass spectrometric approach to analysis, unlabelled lipids generated by the cell (now referred to as the *de novo*[dn]-lipidome [Fig. 1A-B blues]) and lipids incorporating the isotope-labels from FA uptake (now referred to as the *uptake*[up]-lipidome [Fig. 1A-B yellows]) were easily distinguishable by their mass (example structures in Fig. 1). Comparison of relative GPL abundance (Fig. 1B) is preferred over absolute abundance (Fig. 1A) as this minimized uncertainty when comparing between experiments. This comparison revealed that saturated FA supplementation had no apparent impact on GPL class populations (Fig. 1B) although slight differences in contributions of *de novo* (blue) and *uptake* (yellow) fractions from either 13PA or 13SA cells could be observed. In contrast, much greater variation was observed when exploring abundance shifts of the top 15 PC sum composition lipids (Fig. 1C-D), including specific differences between the *de novo*- and *uptake*-lipidome(s). For example, statistically significant increases in unlabelled PC 34:0 within the *de novo*-lipidome were apparent after the labelled 13PA (p≤ 0.01) and 13SA (p≤ 0.05) supplements were introduced (Fig. 1D blues). Concurrently, within the *uptake-*lipidome (Fig. 1D yellows), 13PA or 13SA caused significant increases in PUFA PC lipids, PC 34:2 (p≤ 0.01 & p≤ 0.001), PC 34:3 (p≤ 0.001 & p≤ 0.05), and PC 36:3 (p≤ 0.05 & p≤ 0.05).

**Fig. 1:**
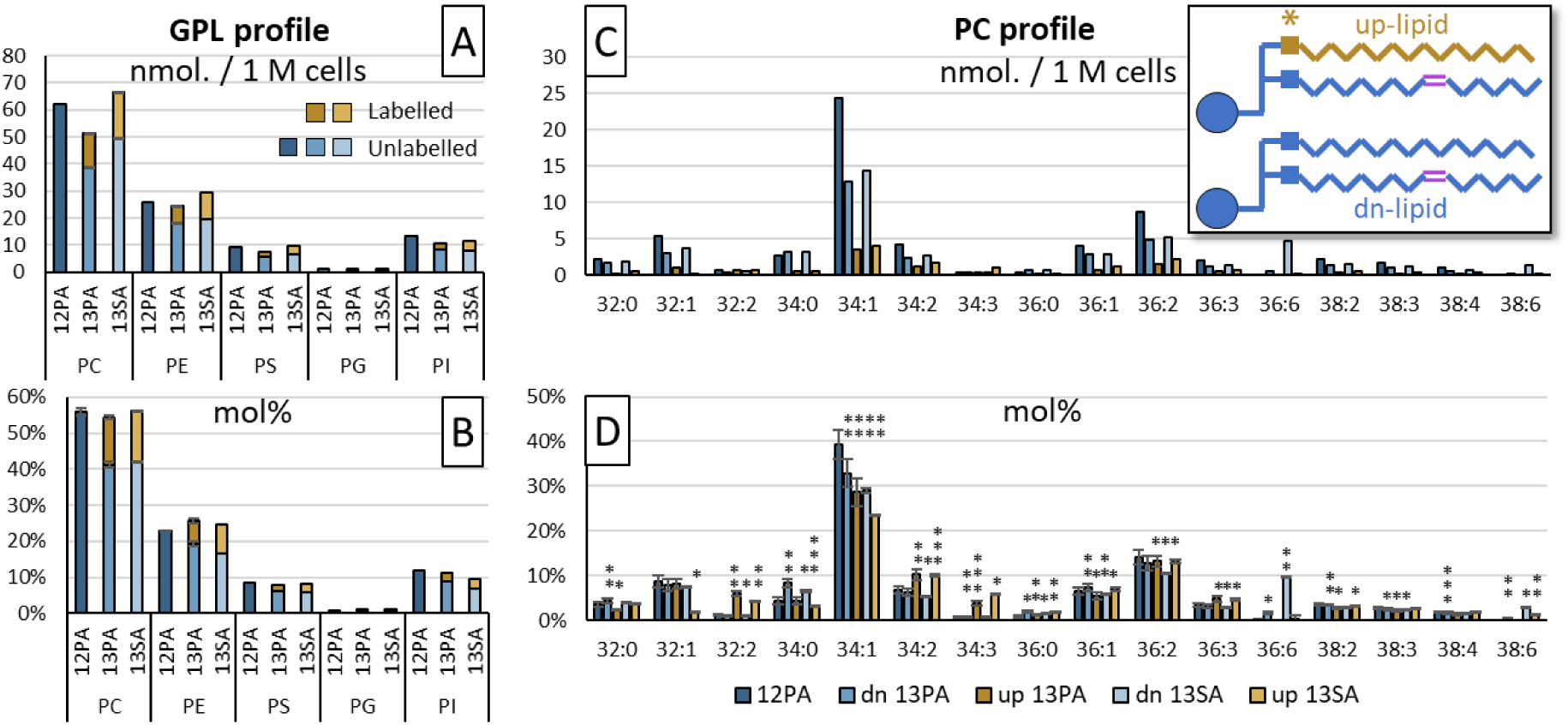
The abundance of glycerophospholipids classes and PC sum compositions (including label-incorporated species) arising from 12PA, 13PA and 13SA supplementation. **(Left)** GPL sum compositions summed by class and separated by *de novo*-lipidome (blue) and *uptake*-lipidome (yellow) abundances; represented in **(A)** absolute abundance (nmol. / 1 M cells), and **(B)** relative abundance (mol%). **(Right)** The effects of supplementation on the top 15 most abundant PC lipids; represented in **(C)** absolute abundance (nmol. / 1 M cells), and **(D)** relative abundance (mol%). Biological replicates: n=2 (NS: n=3) and technical replicates: n=7, mean ± 95% confidence interval displayed. Paired t-test p-values: *0.05, **0.01, ***0.001. Abbreviations: ^12^C_16_-palmitic acid supplement (12PA), ^13^C_16_-palmitic acid supplement (13PA), ^13^C_18_-stearic acid supplement (13SA), *de novo*-lipidome (dn), *uptake*-lipidome (up).

Considering the variation between the *de novo*- and *uptake*-lipidome(s) at the sum composition level, *sn*-isomer analysis populations were established using CID/OzID. Observing the *sn*-isomer compositions of PC 32:1 within the *de novo*-lipidome (Fig. 2A), LNCaP cells grown without FA supplementation (NS) were seen to favour canonical *sn*-2 unsaturated lipids, showing a combined apocromer contribution of 13% the total *de novo-*PC 32:1 (9% PC 16:1/16:0 [light orange] and 4% PC 18:1/14:0 [light grey]). Supplementing LNCaP cells with 12PA, 13PA or 13SA was decreased the overall contribution of canonomers PC 14:0/18:1 (3-5%) and PC 16:0/16:1 (3-5%), resulting in a concomitant increase of the apocromer PC 16:1/16:0 (8-10%). Fig. 2B shows the individual percentage contribution of canonomer (white) and apocromer (black) for the two FA compositions comprising *de novo*-PC 32:1. Notably, this showed that the uptake (but not incorporation) of extracellular FA increased the relative abundance of apocromer species in the *de novo*-lipidome. As with changes to FA *sn*-position, variation in the distribution of DB locational isomers within each supplement experiment was observed (Fig. 2A pie charts). Comparative to NS cells, FA supplemented cells (12PA, 13PA, 13SA) exhibited slight decreases of *n*-8 (≤ 1%), *n*-9 (1-3%) and *n*-10 (1-2%) and a concomitant increase of *n*-7 (3-5%). Together, this combination of increased apocromer contribution (Fig. 2A bar charts) and increased *n*-7 DB isomers (Fig. 2A pie charts) suggested an association between the *n*-7 DB and the *sn*-1 position of PC 32:1. A similar pattern was evident in the PC 34:1 analysis (Fig. S1D pie charts), where *n*-7 and *n*-10 appeared to be positively correlated with sn-1 isomers, while *n*-9 correlated with *sn-*2.

**Fig. 2:**
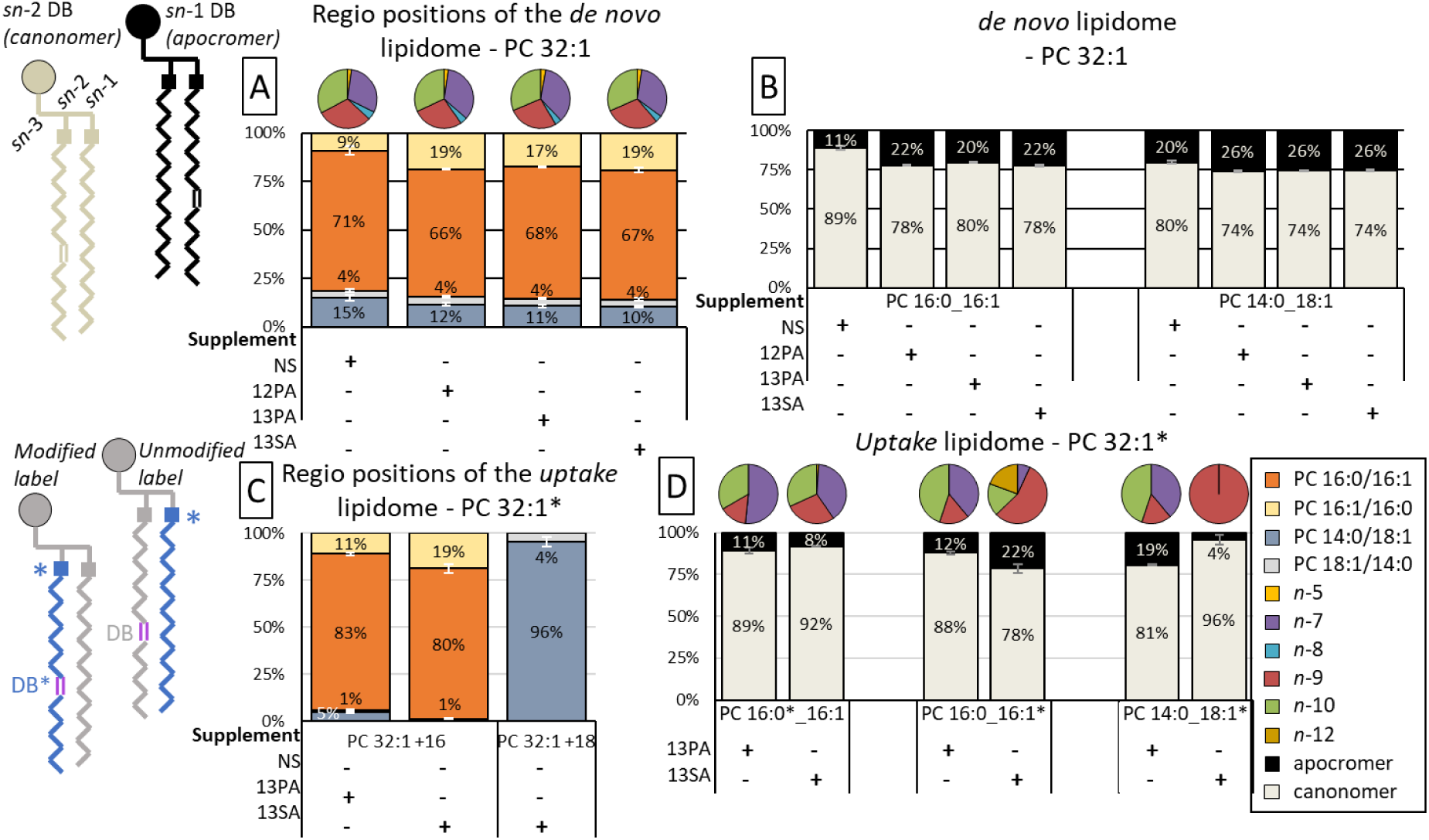
The distribution of DB and *sn-*isomers within the de novo and uptake lipidomes after fatty acid supplementation. **(A)** Bar charts showing the FA *sn-*isomer distribution of two *de novo* PC 32:1 FA compositional isomers, namely, PC 16:0_16:1 (oranges) and PC 14:0_18:1 (greys) and the pie charts showing the distribution of DB isomers within these two lipid species. (**B)** Proportions of PC 32:1 apocromer (black) and canonomer (cream) for each cell supplement - representations of the chemical structure for each are found in the top left. **(C)** Bar charts showing the *sn*-isomer distribution of two *uptake* PC 32:1 lipid species, namely, PC 16:0_16:1 (oranges) and PC 14:0_18:1 (greys). **(D)** Labelled PC 32:1 lipid displayed in panel C can either have the labelled FA unmodified or modified by desaturation and/or β-oxidation (chemical structures represented in the bottom left). The proportions of apocromer (black) and canonomer (cream) are compared between these unique isomers and isotopomers with the labelled FA chain being indicated with an asterisk. Pie charts in (D) are representative of the double bond isomer distribution from the unsaturated FA chain. The values displayed throughout are the relative abundance mean for biological replicates: n=2 (NS: n=3) and technical replicates: n=7. FA supplements are indicated below each chart (+) and isotope labelling is indicated with an asterisk (*).

Investigating the *sn-*isomer distributions of the isotope labelled GPLs that constituted the *uptake*-lipidome of 13PA and 13SA cells, the labelled FAs from uptake were observed to either be directly incorporated into the lipid or were incorporated in combination with cellular modification (*e*.*g*., desaturation, elongation, or β-oxidation). It has previously been shown that the ordering of these metabolic events determines the species of the FA.^25^ The strength of supplementing with isotopically labelled FAs combined with mass spectrometric fragmentation strategies for isomer elucidation is that the shift in *m/z* caused by the inclusion of isotopes provides additional confirmation of FA modification processes. For example, using CID and CID/OzID strategies, the 13SA supplement (*i*.*e*., FA 18:0) displayed four distinct metabolic fates prior to (or post) incorporation into PC 32:1 (Fig. 2C-D pie charts). These included: (i) direct desaturation to form FA 18:1*n*-9, (ii) β-oxidation to form FA 16:0, (ii) sequential β-oxidation and desaturation to form FA 16:1*n*-7 and FA 16:1*n*-10, and the inverse sequence (iv) desaturation and β-oxidation to form FA 16:1*n*-9. Additionally, FA 16:1*n*-12 was formed, which given literature on desaturase Δ-position specificity, could potentially arise from either sequence of β-oxidation and desaturation or direct Δ4-desaturation of FA 16:0. Notably, the modification sequence that the labelled FA underwent was observed to influence the *sn-*position at which it was incorporated (Fig. 2C-D bar charts). For example, compared to the s*n*-positional isomer distribution of NS LNCaP (Fig. 2B), in Fig. 2D the β-oxidation of 13SA to FA 16:0 increased favourability of the canonomer fraction, while combined β-oxidation and desaturation of 13SA to FA 16:1 increased favourability of the apocromer fraction.

As was expected, 13PA cells exhibited an increase within the relative abundance of the PC 16:0_16:1 composition (Fig. 2C). Unexpectedly however, 13SA cells exhibited a greater increase of PC 16:0_16:1 (β-oxidation) over PC 14:0_18:1 (direct desaturation). For comparison back to the *unlabelled* lipidome (Fig. 2B), the *labelled* (*) PC 32:1* data was investigated as a percentage contribution of canonomer or apocromer species (Fig. 2D). As was observed for the 13PA cells in the left and mid panels of Fig. 2D, the 13PA supplement and its modified progeny (FA 16:1*) favour the apocromer (PC 16:1/16:0) with an affinity of 11-12% – equivalent to the isomer distribution of NS cells (Fig. 2B left; 11%). Within 13SA cells, for the 13SA supplement (FA 18:0*) to incorporate into PC 16:0_16:1 it required cellular modification by β-oxidation (to form FA 16:0*) or by combined β-oxidation and desaturation (to form FA 16:1*). When FA 16:0* was formed from 13SA, it showed an 8% affinity for the *sn*-2 position (PC 16:1_16:0*; Fig. 2D left) – ∼3% less favoured than NS cells (Fig. 2B left; 11%). In contrast, when 13SA cells formed FA 16:1* (Fig. 2D mid), this instead occupied the *sn*-1 position (PC 16:1*/16:0) in ∼22% of the species, which was similar to the unlabelled*/de novo* lipidome of cells having received FA supplements (Fig. 2B left; 20-22%). Shifting focus to the variation observed within the PC 14:0_18:1 FA composition, when the 13PA supplement from 13PA cells underwent two cellular modification events (*i*.*e*., desaturation and elongation; Fig. 2D right), FA ^13^C16-18:1* displayed an affinity of ∼19% for the *sn*-1 position (PC 18:1*/14:0). This result is near-identical to the *sn*-isomer distribution of PC 14:0_18:1 within NS cells (PC 18:1/14:0; 20%; Fig. 2B right). 13SA cells again showed contrast with this result, with modification of the 13SA supplement by direct desaturation showing a 4% affinity of the FA 18:1* to the *sn*-1 position (PC 18:1*/14:0; Fig. 2D right). Together, these results present that within 13PA cells the incorporation of 13PA and its progeny to *labelled* PC 32:1* showed close correlation with the *sn*-isomeric distribution of the *unlabelled* PC 32:1 within NS cells. In contrast, within 13SA cells, the incorporation of the cell-modified 13SA supplement to specific *sn*-positions of *labelled* PC 32:1* was significantly influenced by the types and number of modifications it underwent.

Considering that the fatty acyl composition (and *sn-*isomer distribution) of *labelled* PC 32:1* have been established (Fig. 2D), double bond (DB) isomers can be associated with specific lipid species, as is seen within the pie charts of Fig. 2D. Exploring the position of DBs in *labelled* PC 32:1* from cell samples using OzID,^26^ the site of unsaturation was seen to exist in either the unlabelled/*de novo* FA or the labelled/extracellular FA. If the cells were supplemented with 13PA and this remained unmodified (*i*.*e*., unlabelled FA 16:1 undergoes desaturation), *n*-10 fractions remained the same, while *n*-9 is decreased and *n*-7 is increased. If 13PA was provided and this underwent desaturation, *n*-7 and *n*-10 fractions increased, while *n*-9 fractions decreased. From supplementation with 13SA cells, it could be observed that *n*-9 was the only detectable DB position in the label when being incorporated to PC 14:0_18:1*. Interestingly, if 13SA underwent both β-oxidation and desaturation (FA 16:1*), an additional FA 16:1*n*-12 DB isomer arose alongside the increase in FA 16:1*n*-9 – perhaps indicating the origin of the species from a specific subcellular compartment. Alongside the *sn*-isomer distributions (Fig. 2D bar chart), these DB isomer fractions (Fig. 2D pie charts) suggested that *n*-7 FAs are more likely associated with the *sn*-1 position and that FA 18:1*n*-9 has a high affinity to the *sn*-2 position.

#### The relationship between *sn*- and DB position

To investigate an explicit relationship between DB and *sn-*positional isomers, mass spectral dimensionality was increased (MS^4^) to combine both OzID and CID/OzID experiments (*i*.*e*., ordered as OzID/CID/OzID). As such, the distribution of canonomer and apocromer for each FA DB location is quantitative, however absolute mole quantitation of each DB FA sn-isomer would still require establishing OzID reaction rates or developing calibration curves. This technique is not without limitations, however; to achieve the analysis requires: (i) the lipid to be detectable in positive polarity MS, (ii) the precursor ions ability to form sodiated adducts, and (iii) high concentrations of ozone to be generated and passivated into the high-pressure trapping regions of the MS instrument. As such, applications of this technique are currently limited to PC lipids with high cellular abundance such as PC 32:1 and PC 34:1.

Applying this methodology to PC 32:1 from NS LNCaP (Fig. 3) revealed the presence of two fatty acyl composition isomers (PC 16:0_16:1 and PC 14:0_18:1) with the DB in each existing in one of three possible locations (*n*-7, *n*-9, or *n*-10), and with the monosaturated FA being in either canonomeric (*sn*-2) or apocromeric (*sn*-1) positions on the glycerol backbone. In total, 12 discrete isomers were identified within the PC 32:1 sum composition from NS LNCaP. From a quantitative perspective, within the PC 16:0_16:1 composition (Fig. 3 left) the *n*-7 (purples), *n*-9 (reds) and *n*-10 (greens) DBs appear in the apocromer PC 16:1/16:0 (light colours) with a frequency of 29% (±4), 46% (±9) and 54% (±3), respectively. Within the PC 14:0_18:1 composition (Fig. 3 right), the *n*-7, *n*-9 and *n*-10 DB isomers displayed an *sn*-isomer distribution that was distinct from PC 16:0_16:1, with apocromer PC 16:1/16:0 contributing 63% (±11), 27% (±8) and 68% (±5), respectively. These results supported the findings from the aforementioned analyses and confirmed that the 18:1*n*-7 isomer indeed had a higher probability of existing in the *sn*-1 position of the PC 14:0_18:1 species (*i*.*e*., PC 18:1*n*-7/14:0). Similarly, *n*-10 DBs display a similar preference for the *sn*-1 positions of the PC 16:0_16:1 and PC 14:0_18:1 species (*i*.*e*., PC 16:1*n-*10/16:0 and PC 18:1*n*-10/14:0), while *n*-9 shows mixed *sn*-position occupancy that appeared dependant on fatty acyl composition (*i*.*e*., increased relative abundance of both PC 16:1*n-*9/16:0 and PC 14:0/18:1*n*-9 isomers). An equivalent full structure analysis for PC 34:1 can be found in Fig. S1E and showed that within the PC 16:0_18:1 species, *n*-7 and *n*-10 DBs were associated with *sn*-1 isomers, while *n*-9 showed strong preference for the *sn*-2 position.

**Fig. 3:**
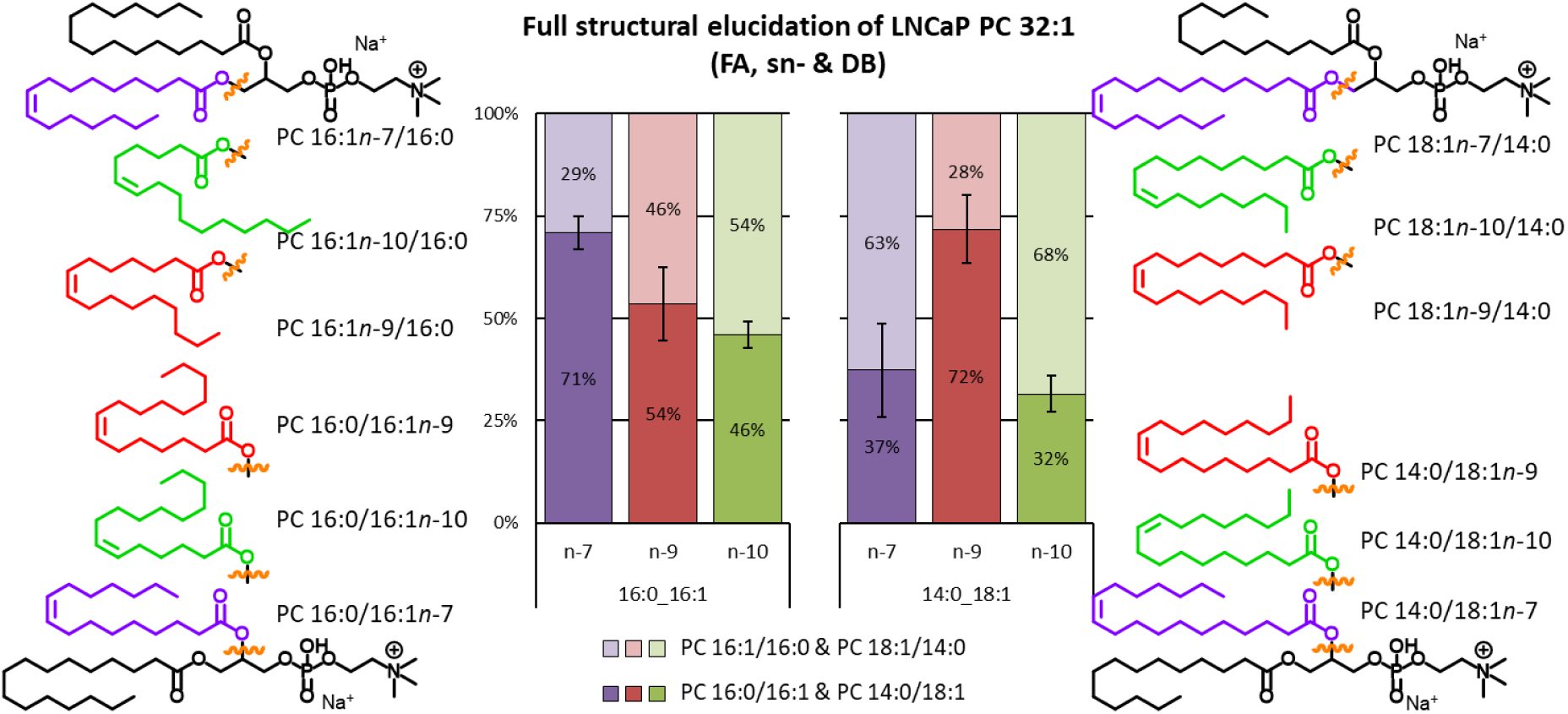
Simultaneous elucidation of fatty acyl composition, *sn-*isomeric position, and double bond locations for the full structural elucidation of PC 32:1 within LNCaP cells. Bar charts showing the *sn-*isomeric distribution (apocromeric in light and canonomeric in dark colours), for each of three double bond positions (*n*-7: purple, *n*-9: red, *n*-10: green) across the two fatty acyl compositional isomers (PC 16:0_16:1, PC 14:0_18:1) within the PC 32:1 species. The 12 molecular structures for each of the measured species are located either side of the chart. n=3, mean value ± SEM (95% confidence interval).

### 3.3 Influence of saturated FA supplements on unsaturated membrane lipids

Fig. 4 displays the widespread effects FA uptake has on GPL lipid remodelling and FA degree of unsaturation. Complex lipidomic analysis across five GPL subclasses was conducted using the 12PA, 13PA and 13SA cell samples. Within all GPL classes the uptake of saturated FAs impacted the degree of unsaturation in both the unlabelled/*de novo-* and labelled/*uptake-* lipidomes (for full lipid profiles see Fig. S2). Using the PS class as an example (Fig. 4B), lipids were separated by the degree of unsaturation to reflect the total contribution of saturated (SFA; blue), monounsaturated (MUFA; orange) and PUFAs to PS profiles. PUFA PS was further separated into lipids with 2 or 3 DBs (as a proxy for mainly *de novo* desaturation; green) and lipids with 4, 5 or 6 DBs (to approximately represent lipids containing “dietary” FAs likely obtained from uptake; yellow). In Fig. 4B and relative to the 12PA cells, PS within the unlabelled *de novo-*lipidome (dn) of 13PA cells appeared to display similar fractions of SFA, MUFA and PUFA. Notably, the ordering of these mole fractions was maintained between 12PA and dn-13PA – with MUFA being the most abundant, sequentially followed by PUFA ≤ 3, PUFA ≥4 and SFA. In contrast and again relative to the 12PA PS profile, the PS within the labelled *uptake-*lipidome (up) of 13PA cells displayed an increase of SFAs along with decreases in MUFA and PUFA ≥4. Interestingly, the degree of unsaturation ordering these lipids within up-13PA is perturbed and instead showed PUFA ≤ 3 being almost equally abundant as MUFA, and SFA now being more abundant than PUFA ≥4. When comparing the PS lipids of up-13SA, slight increases within the PUFA ≥4 fraction (5%) were observed, however the ordering of degree of unsaturation fractions appeared equivalent to the PS profile from 12PA. When instead comparing the dn-13SA PS profile, largescale change could be observed, with MUFA, PUFA ≤ 3 and PUFA ≥4 displaying approximately equal abundance. What is most intriguing about this perturbance in the PS lipids of dn-13SA is that the 13SA supplement itself did not contribute to the observed variation as this lipidome fraction did not carry any isotopic label. Instead, this result implies that stearic acid from uptake induced lipid remodelling to include twice as much PUFA ≥4 to the unlabelled *de novo*-PS fractions, while this effect is only minor in any newly formed PS that incorporated the 13SA label in the labelled *uptake-*lipidome. This effect was apparent within all monitored GPLs (*cf*. Fig. S2).

**Fig. 4:**
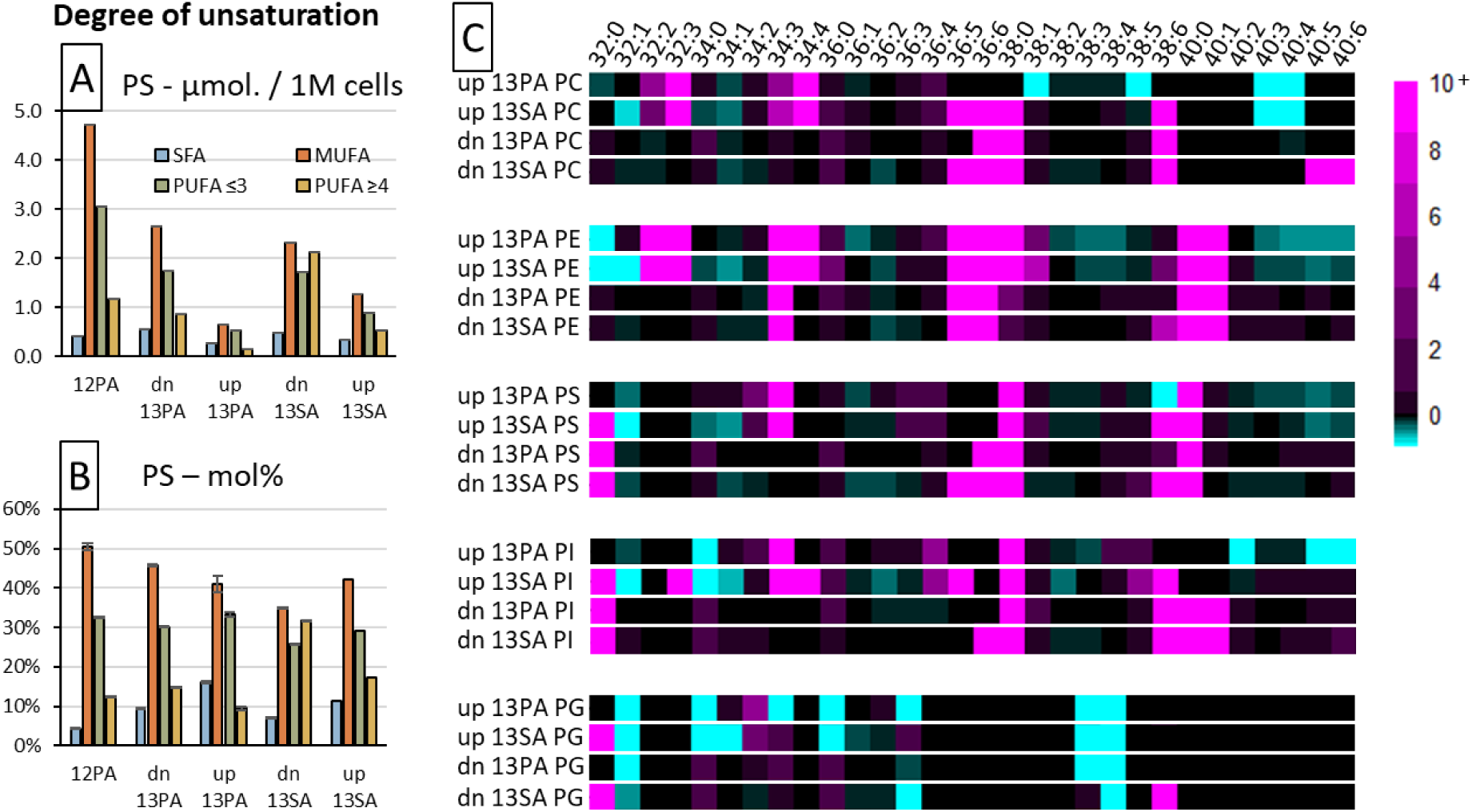
The effects of FA supplementation on the degree of unsaturation of glycerophospholipid species. **(Left)** Bar charts showing the degree of unsaturation in phosphatidylserines in either absolute (A) or relative (B) abundance scales. Degree of unsaturation has been separated into saturated (SFA - blue), monounsaturated (MUFA - orange), polyunsaturated with 2 or 3 DBs (PUFA≤ 3 - green) or polyunsaturated with 4,5 or DBs (PUFA≥4 - yellow). **(C)** Heatmap showing the influence of 13PA or 13SA fatty acid supplementation on either the labelled/*uptake-* (up) or unlabelled/*de novo-* (dn) lipidomes relative to the lipidome arising from 12PA supplementation. Sum composition species are indicated across 5 glycerophospholipid classes. Biological replicates: n=2 and technical replicates: n=7, mean value displayed for absolute abundance or mean value ± 95% confidence interval displayed for relative abundance.

Considering the independent changes to the degree of unsaturation within the *de novo-* and *uptake-*lipidomes after FA supplementation, it was decided that individual lipid speciation should be explored for any variation. To achieve this, a heat map was created where the mol% of FA sum-compositions across five GPLs from the *de novo-* and *uptake-*lipidomes of 13PA and 13SA cells were compared against 12PA cells as a fold-change (Fig. 4C). Overall, this effect of increased PUFA for dn-13SA could be observed across all GPL classes and was exemplified by increases in PI, PS, PC and PE 36:5 and 36:6, and also PI, PS, PC, PG and PE 38:6. Although affecting fewer GPL classes, the up-13SA profiles also displayed this effect but additionally showed decreases in PE 38:4 and PE 40:4 and increases in PC, PE and PI 34:4. The change within GPLs with 5 or 6 degrees of unsaturation (*i*.*e*., EPA or DHA) was less significant in both dn- and up-13PA profiles, however the impact 13PA supplementation had on GPLs with 4 degrees of unsaturation (*i*.*e*., AA) was more widespread than 13SA.

As mentioned previously, PUFA ≥4 is broadly representative of “dietary” FAs, and its increase amongst all monitored dn-GPLs after PA or SA supplementation suggests that these PUFAs are being sourced and remodelled from another lipid class, such as other GPLs or more likely, being sequestered from triacylglycerols (TG).^27-28^ Interestingly, within the dn-13SA profiles an increase within the PC 40:5, PC 40:6 and PE, PS, PI 40:6 could be observed. This is in contrast to the up-13SA profiles, where PS, and PE, 40:5 and 40:6 were seen to decrease. Together these increases in 40:5 and 40:6 from the *de novo*-lipidome and decreases in PUFA PS and PE from the *uptake-*lipidome suggest that the remodelling of PUFA from other lipid classes after FA uptake was GPL class specific. Overall, these results display that the uptake of FAs not only impacted the pre-existing (*de novo*) lipidome but was also influential in the remodelling of PUFAs between the GPLs (and other unmonitored classes) in a class-specific manner.

### 3.4 Implications of FA signalling and lipid remodelling in tissue

We have shown that FA supplementation influences the position and degree of unsaturation as well as the *sn*-positions of FA in PCa cells. These nutrient-driven changes are highly relevant to *in vivo* type experiments, such as xenograft tissue models, where extant cancer cells are frequently subject to variations in the physico-chemical environment and nutrient supply. Indeed, these environmental variations also better represent the usual conditions that cancer cells regularly must endure and thus improve applicability of findings to human cancers. To search LNCaP xenograft tumours for the signatures of metabolic shifts discussed earlier, we deployed mass spectrometry imaging technologies capable of lipid-isomer resolution (MALDI-MSI-OzID).

Fig. 5 shows the MS signal intensity for lipid species as a function of position to display the relative abundance of the indicated lipid across the resected LNCaP xenograft tumours. Sum composition lipids (Fig. 5A-F) were characterised with high mass-resolution and subsequently identified by comparisons to theoretical exact mass (≤ 3 ppm), and GPL *sn-*isomers were resolved using CID/OzID. To compare and correlate the distributions of these lipids (Fig. 5G-I), fractional distribution images (FDI) were created.^21, 29-30^ All images were compared against the structural features of the adjacent H&E stained tissue (Fig. 5J). The H&E image reveals two main features; a round tumour bolus that is dense with cancer cells, and host adipocyte cells surrounding the bolus. Comparing the sum composition lipid images of the top row to the tissue H&E stain, particular lipids showed high levels of spatial correlation with either of the tissue features. For example, PC 32:0 (Fig. 5A) appeared in higher abundance at the perimeter of the fatty tissue, while in contrast PC 36:1 (Fig. 5C) was confined to the centre. It has previously been shown that the acyl chain composition of these two species is dominated by the 16:0_16:0 and 18:0_18:1 fatty acyl compositions,^31^ which represent two unique pathways for formation. It is interesting to see that these two unique synthesis pathways are also spatially distinct, which alongside the differences observed in the labelled supplement study, might suggest that the production and release of palmitic and stearic acids differs between the tumour cells and host adipocyte cells.

**Fig. 5:**
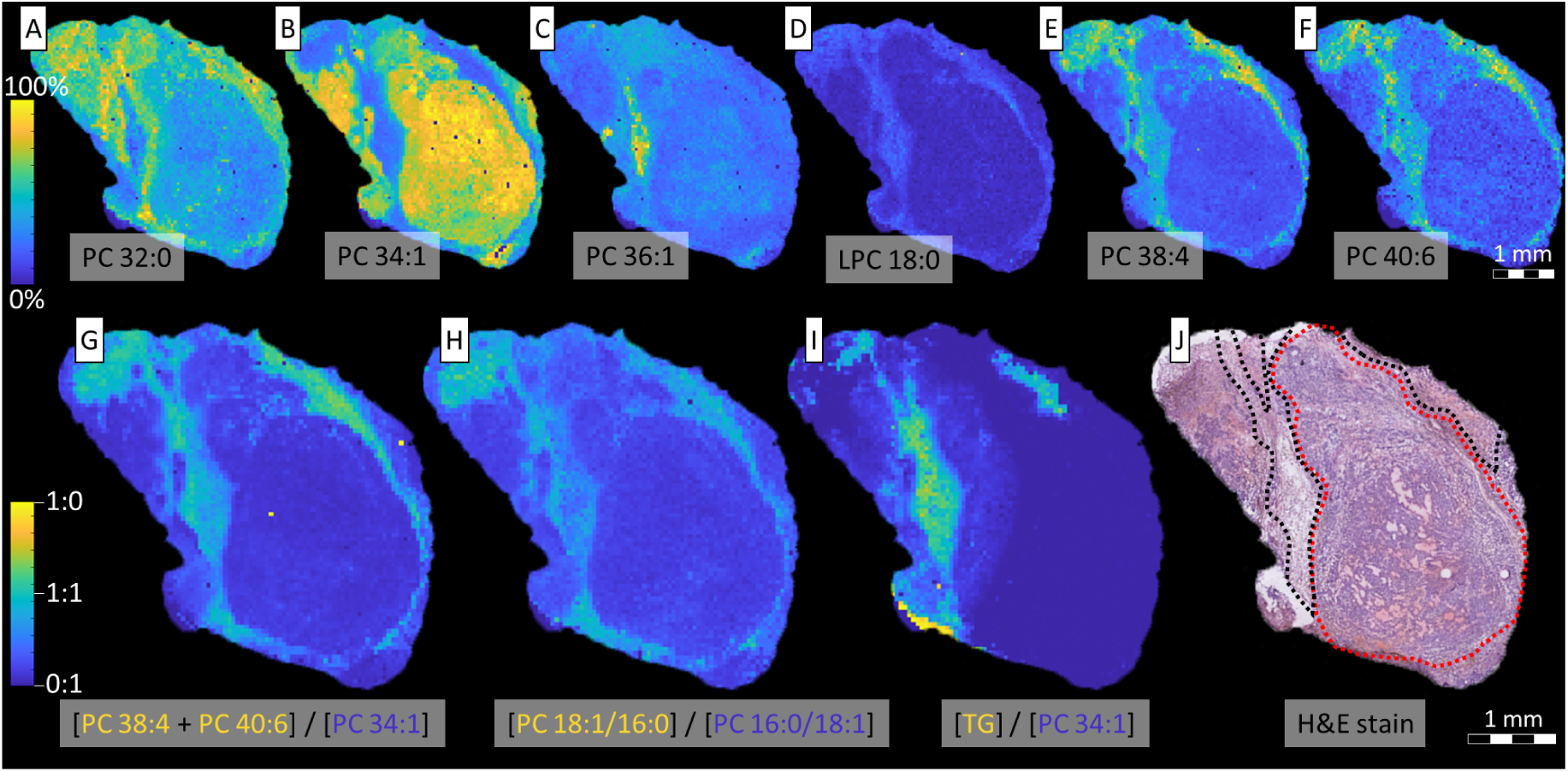
MALDI-MSI and MALDI-MSI-OzID molecular imaging of lipids and the H&E stain of resected LNCaP xenograft tissues. (A-F): The abundance and spatial distribution of (A) PC 32:0, (B) PC 34:1, (C) PC 36:1, (D) lyso-PC (LPC) 18:0, (E) PC 38:4 and (F) PC 40:6. (G-I) Tissue FDIs expressing the relative abundance of the indicated lipid(s) as a function of both lipids for (G) [PC 38:4 + PC 40:6] / PC 34:1, (H) [PC 18:1/16:0] / PC 16:0/18:1, and (I) [TG (48:0, 48:1, 48:2, 50:0, 50:1, 50:2, 50:3, 50:4, 52:0, 52:1, 52:2, 52:3, 52:4, 52:5, 52:6, 54:1, 54:2, 54:3, 54:4, 54:5, 54:6)] / PC 34:1. (J): The H&E-stained tissue showing the tumour cells (red) and host adipocyte cells (black) in a representative treatment-naïve (vehicle) LNCaP xenograft.

One of the more striking observations is the large variation in the abundance of PC 34:1, in either the tumour or host adipocyte cells. While this particular lipid appears to be more abundant in the bolus and less so in the fatty tissue, other lipids such as LPC 18:0, PC 38:4 and PC 40:6 display the complete inverse. This was best observed within the FDI image (Fig. 5G) which revealed that PUFA-GPLs, PC 38:4 and PC 40:6, were indeed spatially distinct to high-abundance PC 34:1. If PC 34:1 content was divided into its *sn-* positional isomer distributions (*i*.*e*., PC 16:0/18:1 and PC 18:1/16:0; Fig. 5H) and compared against the spatial distribution of PUFA-GPLs (Fig. 5G-H), the apocromer showed high positive correlation with the presence of these species. Given that PUFA-GPLs (*e*.*g*., PC 18:0_20:4 and PC 18:0_22:6) are the primary substrate for PLA2,^7^ and the intermediary products of this activity are lyso-lipids, the presence of a lyso-PC (LPC) 18:0 (Fig. 5D) in this same region is suggestive of PLA2 activity. With the altered pH environments that are commonplace in tumours,^22^ acyl chain migration^24^ occurring in these lyso-lipids prior to lipid-remodelling and the reformation of diacyl-lipids would lead to the occurrence of *sn-* apocromer lipid species. Indeed, the overexpression of group IIA secretory-PLA2 (sPLA2-IIA) is observed in LNCaP cells and is correlated with poor clinical prognosis in prostate cancers.^32^ Within murine models that were treated with an sPLA2-IIA inhibitor (KH064), correlations between PUFA GPLs and apocromeric lipids were less apparent, suggesting that this migration activity can be partially attenuated by limiting lipase activity (*cf*. Fig. S3).

An unexpected result (Fig. 5I) is the correlation between TG species and the region of extensive lipid remodelling in the host adipocyte cells. Recent studies investigating how external stimuli can affect cancer cell lipid homeostasis found that the degree of FA unsaturation in TG and GPLs was disrupted by nutrient deprivation and hypoxia.^27, 33^ Given that the physical conditions in the study attenuate cellular FA desaturation, researchers were able to show that in order to maintain appropriate ratios of membrane-bound saturated and unsaturated FAs, cells sequestered unsaturated FAs from TG fractions and transferred to membrane GPLs. Therefore, due to the colocalization of TG and PUFA-GPLs, the increased abundance of PUFA species in GPLs is assumed to come from the repartitioning of TG stores. Similar to the TG localisation, the fatty tissue appears to house an increased abundance of PUFA ether-lipid species (PC O-32:0 and PC O-34:1) and the tentatively assigned TG ethers (*cf*. Fig. S4). These have been previously identified in mammalian lipid droplets and play a central role in ether lipid metabolism and intracellular lipid traffic.^34^ Speculatively, given that the main route for DHA synthesis requires peroxisomal β-oxidation^35^ and the peroxisomes are fundamental in the production of ether lipids,^36^ this data suggests differences in peroxisome metabolism between the host adipocyte cells and tumour cells. Indeed it was recently found that tumour cells can evade ferroptosis by downregulating the peroxisomal synthesis of PUFA ether lipids.^37^

## 4.0 Discussion

With the exception of mitochondria, FA modification is predominantly undertaken in the endoplasmic reticulum (ER) and thus trafficking of FAs and lipids is necessary to reach the plasma and organelle membranes. Although there are exceptions, lipids mainly require active transport mechanisms to shuttle around the cell. The three conventionally accepted trafficking mechanisms involve: membrane transport (*e*.*g*., budding from a donor membrane), carrier proteins (*e*.*g*., fatty acyl binding proteins [FABP]), or transfer across membrane contact sites (*e*.*g*., ER/mitochondria lipid exchange).^38^ Alongside *de novo* lipogenesis, cells can also obtain extracellular FAs from uptake and incorporate these into lipid structures. From our work presented herein, these differential trafficking mechanisms can be inferred from the variations in the PC (and other GPL) lipid species profile(s) (*vide supra* Fig. 1D; for other GPLs *cf*. Fig. S2). For example, tracing the incorporation and modification(s) of labelled palmitic and stearic acids reveals that each FA is either incorporated into more highly unsaturated GPLs or are themselves desaturated more frequently (Fig. 1D; *up* 13PA/*up* 13SA; *e*.*g*., PC 32:2, PC 34:2, PC 34:3 and PC 36:3). In contrast, GPLs solely incorporating FAs synthesised by cellular machinery appear to be influenced by the supplemented FA such that unlabelled PC 34:0 and PC 36:6 species are discretely increased (Fig. 1D; *dn* 13PA/*dn* 13SA). These differences suggest that the cellular management of FAs is dependent both on FA source-origin and FA speciation. Thus we propose that there are distinct mechanisms for the trafficking of FAs originating from *de novo* lipogenesis and those the cell has acquired from uptake. Previous research has also revealed FA-dependent trafficking differences. One such example can be found in FABP5, a FA binding protein that delivers FAs from the cytosol to nuclear peroxisome proliferator-activated receptor beta or delta (PPARβ/δ). Researchers were able to show that while FAs from uptake all complexed with FABP5, a saturated FA (palmitic acid) inhibited FABP5 activation of the PPARβ/δ pathway, while a polyunsaturated FA (linoleic acid; LA; FA 18:2*n*-6) instead activated PPARβ/δ.^39^ This FA-driven inhibition/activation of PPARβ/δ has downstream effects in the activation of lipometabolic-centred genes such as pyruvate dehydrogenase kinase 4 (*PDK4*), angiopoietin 4 (*ANGPTL4*), perilipin 2 (*Plin2*) and cluster of differentiation/FA translocase (*CD36*). Considering the changes that occur to FA metabolism in cancer, especially including FA uptake and energy production from FAs, the translation of these genes into proteins has widespread implications for oncometabolism.^40-45^ Furthermore, this FABP5/PPARβ/δ mechanism highlights how FA trafficking mechanisms can be entangled with signalling events, while this work provides the foundation for the discovery of the proteins involved in partitioning of dn- and up-FAs.

Given the widespread impact FA trafficking can have on signalling for gene expression and protein translation, it is logical to assume that the cell has extremely fine control over FA uptake, FA trafficking and regulation of lipid remodelling. This is exemplified in our work by the DB or *sn-*position differences observed between labelled or unlabelled FAs, and possibly relates to the known compartmentalisation of desaturase enzymes. Within the pie charts of Fig. 2D, the percentage of *n*-10 (requiring FADS2 desaturation) is observed to be higher when the labelled palmitic acid is modified by desaturation or by combined desaturation and elongation, while *n-*7 and *n*-9 (requiring SCD-1 desaturation) are seen to be higher within labelled lipids where the unlabelled FA is the unsaturated chain. This would suggest that a higher proportion of FAs from *uptake* are transported to the mitochondria (where FADS2 desaturation occurs)^46^ while *de novo* synthesised FAs are more readily transported to the ER (where SCD-1 desaturation occurs).^47^ Further, this would indicate that FA speciation and origin (*i*.*e*., uptake or *de novo*) both determine subcellular destination as opposed to the conventional theory that all FAs form a common “pool”. One explanation for this compartmentalisation of FA fractions would be for net-positive energy production. Catabolising FAs that have been actively *de novo* synthesised, although sometimes required, is counterproductive to energy production. Instead, trafficking FAs from uptake to β-oxidation sites would provide externally produced energy (and carbon) to the cell. Interestingly, FA uptake influences PPARβ/δ activity and subsequent activation of PDK4, which regulates the conversion of pyruvate and glucose to acetyl-CoA and supresses ferroptosis.^40^ These functions directly implicate compartmentalised FA metabolism as mitochondria are responsible for cellular energy production via β-oxidation and FA synthesis, while the peroxisomes mediate an alternate FA β-oxidation pathway and are largely responsible for the formation of reactive oxidative species and ferroptosis.

A related result that supports the hypothesis of specific FA species compartmentalisation can be found in the unusual lipid DB profile displaying 16:1*n*-12 and increased 16:1*n*-9 (Fig. 2D) after a labelled stearic acid has undergone partial β-oxidation and desaturation to FA 16:1*. As stated, within the cell both the mitochondria and peroxisomes have the capacity for β-oxidation, however peroxisome β-oxidation and stimulation has been shown to have a complex relationship with the genetic expression of *fads*1 and *fads*2 and hence FADS1 and FADS2 enzymes.^48-49^ These enzymes catalyse Δ4, Δ5, Δ6 and Δ8 desaturation^31, 46^ and in conjunction with β-oxidation would lead to the formation of the unusual 16:1*n*-12 species observed within Fig. 2D (mid). This may suggest that the stearic acid supplement is being split between two trafficking pathways: (i) to the ER for SCD-1 desaturation and then membrane-contact transport to the peroxisomes for β-oxidation to yield 16:1*n*-9 and stimulate the peroxisomal influence on FADS2, and (ii) to the mitochondria for FADS2 desaturation and β-oxidation to yield 16:1*n*-12.

Another example of remodelling compartmentalisation can be observed in Fig. 3 (left), where PC 16:0_16:1*n*-9 and PC 16:0_16:1*n*-10 from non-supplemented LNCaP showed equal apocromer/canonomer distribution, while PC 16:0_16:1*n*-7 markedly favoured canonomeric incorporation. This is a remarkable result as both β-oxidation and FADS2 desaturation (leading to FA 16:1*n*-9 and *n*-10, respectively) are mitochondrial processes, while the SCD-1 desaturation forming FA 16:1*n*-7 would occur at the ER. Similarly, the PC 14:0_18:1 in Fig. 3 (right) shows that *sn*-isomer distribution of FA 18:1*n*-7 and *n*-10 is quite similar, while *n*-9 is distinctly different. To form PC 14:0_18:1, both FA 16:1*n*-7 and FA 16:1*n*-10 would require elongation, while FA 18:1*n*-9 would be directly desaturated from FA 18:0. Interestingly, subcellular organelles are observed to have discrete pH ranges with mitochondrial pH being 7.58 for the matrix and 6.88 for the intermembrane space,^50^ ER pH being 7.1 at a resting state^51^ and peroxisome pH varying between 7.4 and 8.1 depending on metabolic activity and cell-type properties.^52^ Considering acyl chain migration (while lipids exist as lyso-species) is a pH dependant equilibrium, the remodelling of lipids under the different pH conditions of subcellular compartments could indeed lead to differences in the *sn*-positional isomer distributions. Therefore, the *sn-* positional isomers may be a further indication of compartmentalised FA modification events. Because FA 16:1*n*-9 has been recently identified as a mediator for an anti-inflammatory response,^53-54^ FA 16:1*n*-7 is known as a lipokine that regulates SCD-1 expression^55^ and FA 16:1*n*-10 has anti-microbial properties,^56^ it is paramount to observe and measure any changes to the *sn-* isomeric composition in order to better understand the enzymes responsible for their catalytic release in metabolic diseases such as cancer.

FAs in their free form have the potential for signalling cellular processes and mechanisms.^18, 39, 57-59^ Along with previous examples (*i*.*e*., FA 16:1*n*-7, FA 16:1*n*-9 & FA 16:1*n*-10), the FAs comprising the bulk of research surrounding signalling behaviours are those from dietary (extracellular) sources. These include: FA 16:1*n*-7(*trans*) and FA 18:1*n*-7(*trans*) from milk products, which stimulate endocannabinoid and noncannabinoid production to in turn impact peroxisome proliferator-activated receptor alpha (PPARα), lipolysis, inflammation and ANGPTL4;^60-61^ LA and AA mainly from plant sources, which are precursory to eicosanoid formation and the pro-inflammation response;^9^ and, alpha-linolenic acid (ALA), eicosapentaenoic acid (EPA) and docosahexaenoic acid (DHA) from mainly marine sources, which mediate an anti-inflammatory response through resolvins, protectins, maresins and the disruption of lipid rafts.^8, 62^ Further, the presence or absence of specific FA species within particular GPL classes and the degree of unsaturation in membrane lipid FAs has been shown to be influential factors in cell signalling.^63-64^ As seen in Fig. 4C, notably our work shows that the uptake of stearic acid drives a sharp increase of DHA into GPLs. Given that these GPL-DHA species are prime targets for PLA2 activity, the increase in these substrates would likely lead to the release of DHA from the lipid and mediate anti-inflammatory signalling cascades. This signal by *uptake* FAs presents a promising target for inducing pro- or anti-inflammatory responses via non-toxic interventions. For example, by pre-exposing the cells to DHA substrates for a period of time and then supplementing with stearic acid. This should repartition DHA into the GPLs, which upon lipase activity would initiate a strong anti-inflammatory effect and provide a non-toxic alternative to conventional anti-inflammatory drug therapies.

Given that FAs from uptake are shown to signal for the remodelling of PUFAs into the GPL lipid fraction (*cf*. Fig. 4), one explanation for this distinction between *de novo* and *uptake* FAs might lie within homeostasis. Within Fig. 2 it is observed that FA uptake causes a large variation to the abundance of lipid apocromers in the *de novo*-lipidome fraction comparative to the nil-supplemented control. In contrast, the supplementation and incorporation of palmitic acid (and its metabolic progeny) into the *uptake*-lipidome is seen to influence the apocromer abundance such that it replicates the nil-supplemented control. In the more biologically relevant instance where the cell is constantly taking up extracellular FAs that are causing the signalling for remodelling events, it is logical to assume the incorporation of these new FAs into lipids would follow a homeostatic cellular template. Accordingly, the more that FAs are taken up, the larger this uptake-lipid fraction becomes until it exists in majority and thus dilutes the lipid fraction that was perturbed by remodelling – thereby restoring homeostasis. Disruption of these mechanisms, such as changes to environmental physicochemical properties, would therefore lead to the incorrect formation of lipid *sn-* isomers and have dire implications on homeostatic metabolism. The spatial correlations observed in Fig. 5 between PUFA lipids, lipid apocromers, lyso-lipids and TGs indicate that there was dynamic lipid remodelling occurring within the fatty tissue of the xenograft model, however this same remodelling is not observed within the bulk of the tumour (round bolus). Here, it is speculated that the hallmark switch to FA metabolism by tumour cells led to the production and secretion of palmitic and stearic acids, while simultaneously creating a more acidic environment for the surrounding tissues and an increased expression of sPLA2-IIa in attempt to decrease inflammation. Subsequently, the secreted palmitic and stearic acids were taken up by the host adipocyte cells, which caused the repartitioning of PUFAs from TGs to GPLs, especially PIs which are mainly located on the outer leaflet of the plasma membrane. These outwardly exposed PUFA PIs are a prime target for the increased presence of sPLA2-IIa, which would in turn lead to the formation of lyso-lipid and free FA species. The hypothetical change in the pH caused by the tumour would then lead to lyso-acyl chain migration and the formation of apocromeric lipid species. Consequently, lipids normally containing signalling FAs at the *sn*-2 position, that would be released by PLA2 under normal circumstances, would now release and excrete the *sn*-2 saturated FA – which could potentially go on to initiate the same cataclysmic remodelling response in surrounding cells. Considering the changes that are occurring within the TG, ether-lipid (TG O-, PC O-) and GPL sum compositional fractions (*cf*. Fig. 5 and Fig. S4) and their roles and biosynthetic origins, it is tempting to speculate that the excretion of these saturated FAs would also lead to the earlier mentioned FABP5 inhibition/activation of PPARβ/δ and its influence on the expression of ANGPTL4, Plin2 and CD36. Each of these enzymes can be inferred within the observed changes. Changes to ANGPTL4 (known for its roles in reactive-oxidative species [ROS] regulation, hypoxia response and anoikis)^41-42^ expression would be assistive in the prevention of ferroptosis brought on by ROS species coming from increased high levels of GPL unsaturation and lipid peroxidation. Interestingly, the formation of ROS species is also controlled by the peroxisomes, which along with previous implications, peroxisomal function/dysfunction can also be implied through the co-distribution of ether-lipid species (peroxisomal lipid class) to the PUFA-GPL tissue regions. TG speciation and spatial distribution (*cf*. Fig. S4) also alludes to changes in lipid droplet formation in PUFA-GPL tissue regions, while Plin2 is known to interact with lipid droplets to indirectly regulate droplet structure and function.^43^ Lastly, changes to FA/lipid uptake and trafficking are readily observed throughout this work, which are known functions of CD36^44^ and Together the relationship between FA metabolism and these downstream effects of FA activated PPARβ/δ activity suggests that the trafficking of FAs and lipids is tightly regulated and highly influential in cell homeostasis.

Although previous research has provided a wealth of knowledge with regards to lipid remodelling, trafficking, and signalling, the recent progress made in lipid isomer research calls for an update on how these processes are impacted by or impact upon isomeric lipid species. Within, we present an array of isomeric changes that occur after FA supplementation and how these mechanisms are manifesting in tumour tissues. Importantly, these metabolic differences are heavily related to known oncogenic enzymes that are influential in cancer pathogenesis. Together, the indication that there exists distinct subcellular fractions of FA that have different metabolic fates, functions and flow-on effects suggests that lipid isomers could be potential biomarkers for lipid remodelling and disease progression in tumour tissues.

## 5.0 Acknowledgements

This work was financially supported by the Australian Government through award of an RTP scholarship (to R.S.E.Y); the Australian Research Council through the Discovery Program (DP190101486) and the Linkage Program (LP180100238, partnered with Waters Corporation); the Prostate Cancer Foundation of Australia and the Australian Government Department of Health through a Movember Revolutionary Team Award; 2019 Cancer Program Initiative for Metabolism (Institute of Health and Biomedical Innovation, QUT); and, the Dutch Province of Limburg as part of the “LINK” program. S.R.E., A.P.B. and R.M.A.H. acknowledge funding from Interreg V EMR and the Netherlands Ministry of Economic Affairs within the “EURLIPIDS” project (project number EMR23). S.R.E. acknowledges funding from the Australian Research Council Future Fellowship Scheme (grant number FT190100082). The project team acknowledge the Central Analytical Research Facility operated by the Queensland University of Technology for provision of instrumentation and training in support of this project; the Translational Research Institute (TRI), including the Biological Resources Facility, Preclinical Imaging and Histology facilities (and supporting grants from the Australian and Queensland Governments); and, David L. Marshall and Gert B. Eijkel for helpful discussions and software developments.

## 6.0 Author contributions

**Reuben S**.**E. Young**: Conceptualisation, Methodology, Software, Validation, Formal analysis, Investigation, Data curation, Writing – Original draft, Writing – Review and Editing, Visualisation, Project administration. **Andrew P. Bowman**: Validation, Investigation. **Kaylyn D. Tousignant**: Validation, Investigation. **Jennifer H Gunter**: Resources. **Lisa K Philp**: Resources. **Berwyck L**.**J. Poad**: Methodology, Resources, Writing – Review and Editing, Supervision, Funding acquisition. **Colleen C. Nelson**: Resources, Funding acquisition. **Shane R. Ellis**: Methodology, Resources, Writing – Review and Editing, Funding acquisition. **Ron M**.**A. Heeren**: Resources, Funding acquisition. **Martin C. Sadowski**: Conceptualisation, Methodology, Validation, Investigation, Writing – Review and Editing, Supervision, Project administration. **Stephen J. Blanksby**: Conceptualisation, Methodology, Resources, Writing – Review and Editing, Supervision, Project administration, Funding acquisition.

## 8.0 Methods

### 8.1 Materials

Methanol (LC-MS grade), acetonitrile (ACN; Optima®), water (Optima®), isopropanol (IPA; Optima® LC-MS grade) for lipid LC-MS were obtained from Fisher Scientific, Scorseby, VIC, Australia. Ammonium acetate for lipid LC-MS was obtained from Sigma-Aldrich, North Ryde, NSW, Australia. For MALDI-MS related sample prep and analysis, sodium acetate (≥99%), 2,5-dihydroxyacetophenone (≥99.5%, Ultra pure), methanol (LC-MS grade) and chloroform (HPLC grade) were obtained from Sigma Aldrich, Zwijndrecht, The Netherlands.

### 8.2 Biological sample preparation

#### 8.2.1 Ethics and murine model

Animal ethics approval was granted by the Office of Research Ethics, University of Queensland and Queensland University of Technology, and assigned AEC approval number QUT/260/18. All testing was undertaken in accordance with accepted standards of humane animal care outlined by the Australian Code of Practice for the Care and Use of Animals for Scientific Purposes. Cell line use was approved by Queensland University of Technology Human Research Ethics. Six-week-old male NOD/SCID mice were purchased from the Animal Resource Centre (ARC, Australia) and maintained under temperature-controlled conditions and a 12 h light-dark cycle. Mice were allowed to acclimate for 2 weeks prior to experiment and were provided chow and water *ad libitum* for the duration of the experiment. Mice were injected subcutaneously in the right flank with LNCaP cells (1 M cells) in a 1:1 ratio of cells and Matrigel® (100 μL; Corning®). Bloods were collected once weekly and prostate specific antigen (PSA) was measured by ELISA to monitor PCa progression. Mice were weighed and tumour volume was measured using digital callipers thrice weekly. Tumours were allowed to grow until PSA reached 25-50 ng/mL before mice were castrated. One week following castration, mice were randomised to receive an sPLA2-IIa inhibitor (KH064; 5 mg^-1^ kg^-1^ day^-1^ by oral gavage) or vehicle control (0.5% Carboxymethylcellulose). Mice were culled when the ethical endpoint was reached, defined as maximum tumour volume (1000 mm^3^) or when an Animal Ethics Committee-approved welfare score required euthanasia. Xenografts and tissues were collected and weighed. Tumours were hematoxylin and eosin (H&E) stained for morphological assessment, while tissues were halved and either: (i) placed in liquid nitrogen until moved to storage at −80°C, or (ii) stored in neutral buffered formalin fixation and paraffin-embedded (FFPE).

#### 8.2.2 Tissue sectioning and mounting

Frozen resected tissues were freeze mounted to the microtome block using water and sectioned at 10 μM thickness for lipid analyses using a CM 1950 Cryostat (Leica Biosystems, Nussloch, Germany), and using a blade that was free from optimal cutting temperature (OCT) compound. Tissue sections were thaw mounted on standard glass slides (SuperFrost Plus, Menzel-Gläser, Braunschweig, Germany) before MALDI-MSI-OzID protocol (see Method 8.3.1) and the procedure used for H&E staining (see Method 8.2.3).

#### 8.2.3 Haematoxylin and Eosin Staining

A standard H&E protocol was used for staining of xenograft tissues (70% ethanol 2x 3 min, water 1x 3min, Hematoxylin 1x 3 min, water 1x 3 min, Eosin 1x 30 s, water 1x 3 min, ethanol 1x 1 min, Xylene 1x 30 s). High-resolution optical images of stained tissues were generated using an Aperio CS2 digital pathology slide scanner (Leica Biosystems, Wetzlar, Germany). Hematoxylin and Entellan® were purchased from Merck (Darmstadt, Germany) and eosin Y from J.T. Baker (Center Valley, USA).

#### 8.2.4 Cell culturing

LNCaP (RRID: CVCL_0395) cells were obtained from the American Type Cell Culture Collection (ATCC; Manassas, Virginia, USA) and were cultured in Roswell Park Memorial Institute (RPMI) medium (Thermo Fisher, Waltham, MA, USA) supplemented with 5% fetal bovine serum (FBS, Invitrogen, Waltham, MA, USA) and incubated at 37 °C in 5% CO2. Medium was changed every 3 days, and cells were passaged at approximately 80% confluency by trypsinisation. Cell lines were authenticated by STR profiling in March 2018 and December 2020 by Genomics Research Centre (Brisbane, Australia) and routinely tested to exclude mycoplasma infection (Lonza MycoAlert). Cell number and viability was determined by trypan blue staining and using a TC20 Automated Cell Counter (Bio-Rad).

#### 8.2.5 ^*13*^*Carbon tracing*

Cells were seeded and cultured under the conditions mentioned previously (Method 8.2.4). After 48 hours of seeding, cells were switched to fresh RPMI media supplemented with 10% charcoal-stripped bovine serum (CSS; Invitrogen, Waltham, MA, USA) and incubated at 37 °C for 30 min to remove FAs from the binding sites of bovine serum albumin (BSA). FA depleted cells were then supplemented with either unlabelled (*i.e*., 12C16-palmitic acid) or labelled (*i.e*., ^13^C16-palmitic acid or ^13^C18-stearic acid) fatty acids at a final concentration of 20 µM. All fatty acids were purchased from Sigma Aldrich, Castle Hill, Australia. Cells were grown for a further 72 hours and washed twice with ice-cold phosphate buffered saline (PBS) before lipid extraction. Cell number and viability were determined by trypan blue exclusion with a TC20 Automated Cell Counter (Bio-Rad).

#### 8.2.6 Lipid extraction

Cell lipids were extracted using methods similar to those described by Matyash *et al*.^65^ and were quantifiable through the use of internal standards in the form of deuterated lipids (SPLASH Lipid-o-mix®, Avanti Polar Lipids, Alabaster, USA). To minimize pipetting error, a stock internal standard solution was made in bulk using 720 µL MTBE (0.01% BHT), 40 µL SPLASH Lipid-o-mix® and 20 µL nonadecanoic acid in MTBE (3.35 mM) per 2 M cells. Cell pellets in 2 mL clear glass vials (∼2 M cells) were twice washed with PBS solution, before adding 220 µL of methanol and 780 µL of the prepared internal standard stock solution. Capped vials were vortexed for 20 s, before 1.5 h bench-top agitation. Phase separation was induced by adding 200 µL of aqueous ammonium acetate (150 mM) before samples were vortexed for 20 s and centrifuged for 5 min at 2,000 x g. The organic supernatant was pipetted off to a clean labelled 2 mL glass vial and stored at -20 °C before analyses.

### 8.2 Analytical methods

#### 8.3.1 Mass spectrometry imaging of lipid isomers

Samples for MALDI-MSI were placed into microcentrifuge tubes, purged with nitrogen gas and stored on dry ice for inter-laboratory shipping. Samples were stored at -80 °C upon arrival. Immediately prior to analysis, tissues were first sectioned as per Method 8.2.2 and thinly coated with 12 passes (45 mm spray height, 30 °C, 10 psi, 2 mm track spacing) of 100 mM sodium acetate dissolved in 2:1 methanol/chloroform using an HTX TM-Sprayer (HTXImaging, Chapel Hill, NC, USA). 2,5-dihydroxyacetophenone was then sublimated (40 mg, 160 °C, 4 mins) to the sample slides using a sublimator (HTXImaging, Chapel Hill, NC, USA). Two separate MALDI-MSI experiments were undertaken on 10 µm tissue sections that were adjacent to H&E stained tissues. The first was the identification and spatial distribution of sum composition lipid species via high mass-resolution (120k at *m/z* 400) mass analysis (≤ 3 Δppm), and the second was the identification and spatial distribution of PC 34:1 *sn-*isomers (using MALDI-MSI-CID/OzID). All MSI experiments were performed on an Orbitrap Elite (Thermo Scientific, Bremen, Germany) mass spectrometer with MALDI-MSI pixel size of ∼50 µm. The commercial ion source and stacked ring ion guide were replaced with an elevated-pressure MALDI ion source incorporating a dual-ion funnel interface (Spectroglyph LLC, Kennewick, USA) as has been described previously.^66^ The MALDI laser was operated at a repetition rate of 100 Hz and pulse energy of ∼1.0 µJ. The laser was focused to a spot size of ∼15×12 µm as determined by the size of ablation craters in desorbed matrix. Pressure within the ion source was set to 10 mbar in the first ion funnel, and 2 mbar in the second ion funnel. The mass spectrometer was additionally modified to allow for ozonolysis, as previously described.^21^ Briefly, using oxygen as a feed gas (99.999% purity, Linde Gas Benelux BV, The Netherlands) ozone was generated by via a high concentration generator (TG-40; Ozone Solutions, Hull, IA, USA) and was introduced into the helium buffer gas flow before conduction through to the high-pressure region of the linear ion trap (LIT). Ozone concentration was measured online using a UV-absorption based ozone monitor (106-H, 2B Technologies, Boulder, CO, USA). For the determination of FA *sn*-position, precursor ions were mass-selected using an isolation width of 1 *m/z*. Operating in positive polarity mode, CID/OzID and full MS scans were alternated between for each pixel. MS^3^ CID/OzID of *m/z* 782 (using NCE of 42 for MS^2^and an AT of 250 ms for MS^3^) was conducted and detected in the LIT and full MS detection was undertaken by the Fourier Transform-orbitrap detector operating with a mass resolving power of ∼60,000.

#### 8.3.2 Direct infusion ESI-OzID of lipid double bonds

The DBs of intact glycerophospholipids were determined via mass spectrometry using an Orbitrap Elite high-resolution mass spectrometer (Thermo Scientific, Bremen, Germany) modified with an ozone generator (Titan-30UHC Absolute Ozone, Edmonton, Canada), as described in Method 8.3.1. In this instance, a diverter valve was placed on the nitrogen gas inlet to the higher collisional dissociation (HCD) cell, and nitrogen was replaced with generated ozone gas.^67^ Operating in positive-ion mode for PC lipid OzID, cell line lipid extract samples were mixed 1:1 (v/v) with 1 mM methanolic sodium acetate solution and introduced to the mass spectrometer via a chip-based nano-electrospray source (TriVersa Nanomate, Advion, Ithaca, NY, USA) using 1.35 kV/0.35 psi spray parameters. Using the Thermo Xcalibur software package, a data independent acquisition sequence was created to perform sequential OzID (activation time (AT) = 5 s (HCD), collision energy (CE) = 1 V) and CID/OzID (MS^2^: AT = 5 ms (LIT), normalized collision energy (NCE) = 33; MS^3^: AT = 500 ms, NCE = 0) for 6 labelled and unlabelled sodiated phosphatidylcholine precursor ion masses. Maximum injection time was 100 ms, isolation window was ± 0.5 Da across a 175-1000 Da scan range. Included sodiated precursor ion m/z values were: 754.6, 782.6, 770.6, 772.6, 798.6, and 800.6. The product ions from all fragmentation experiments were detected using the orbitrap mass analyser for high resolution and accurate mass, allowing for unambiguous assignment of characteristic fragments to specific lipids and not isobaric lipids or isotopes. Intensity values obtained from OzID and CID/OzID mass spectrometric experiments were the average of 11 and 28 scans, respectively.

Adapting the aforementioned positive-ion mode experiments, MS^4^ experiments were created for simultaneous full structure elucidation, including DB location and sn-position. In brief, sodiated PC precursor ions were mass selected to undergo OzID in the linear ion trap (MS^2^: NCE = 0, AT = 10 s). The OzID-aldehyde product ion of one double bond position was then mass selected to undergo CID in the LIT (MS^3^: NCE = 33, AT = 5 ms) to form a product ion corresponding to the loss of phosphocholine, [M-183]^+^. This product ion was then mass selected for additional OzID in the LIT (MS^4^: NCE = 0, AT = 500 ms) to yield 4^th^ generation product ions that are characteristic to FA chain lengths at either *sn*-1 or *sn*-2 positions for the DB location that was selected for MS^3^ activation. Due to the multiple levels of MS for species confirmation, detection was achieved by the low-resolution detector of the linear ion trap. This experiment was then repeated for the remaining OzID-aldehyde product ions from each DB location.

#### 8.3.3 Conventional lipidomics for phospholipid profiles

Lipid extracts from cell lines were run through an automated lipidomics workflow using an LC-20A HPLC (Shimadzu, Kyoto, Japan) set to deliver 100 µL sample loop-injections into a mobile phase of 5 mM methanolic ammonium acetate flowing at 15 µL/min. The sample column and column oven were bypassed with Viper PEEKsil (50 mm, Thermo Fisher, Waltham, MA, USA) to maintain instrument back pressure limits. Sample lipids were then directly infused through the electrospray ionisation source of a QTRAP 6500 hybrid triple quadrupole/LIT mass spectrometer (SCIEX, Concord, ON, Canada) using a spray voltage of 5 kV, a source temperature of 150C and both source gasses set to 15 (arb.). Various precursor ion and neutral loss scans were employed to confirm lipid head group, with the detected m/z being indicative of summed-fatty-acyl composition. (PC: PIS m/z 184.2, CE: 39V; PE: NL m/z 141.1, CE: 29V; PS: NL m/z 185.1, CE: 29V; PG: NL m/z 189.1, CE: 29V; PI: NL m/z 275.1, CE: 29V; ChE: NL m/z 259.1, CE: 29V). Instrument blanks were run through-out to ensure no sample carry-over and pooled batch quality controls (PBQC) were used to gauge instrument performance over the duration of the experiment.

### 8.4 Quantification and statistical analysis

#### 8.4.1 MALDI-MSI OzID for lipid double bond imaging

MS spectral data files were first stitched together with MALDI imaging co-ordinate positions and collated as a MATLAB data table. TIC normalisation of the data was achieved by dividing each spectral peak by the total ion abundance in each MSI pixel and comparing this to all other pixels. Lipid abundance and distribution data were then visualised through MATLAB scripts developed by Gert. B Eijkel.^21, 66^ Functional distribution images (FDI) were calculated by plotting the intensity of one monitored ion divided by the combined intensity of all monitored ions (*i.e*., *Ia* / [*Ia + Ib*])

#### 8.4.2 Conventional lipidomics for phospholipid profiles

Lipidview (Version 1.3 beta, SCIEX, Concord, ON, Canada) was used for data processing of SCIEX data files obtained from the QTRAP 6500. Lipid assignments were based on the software lipid tables and shortlisted to include even-chain lipids with 0-6 s. Odd-chain/ether-lipid data was obtained but was not included in this study due to the ambiguity in assigning isobars in low resolution mass spectrometry. Isotope correction factors were applied, and MS peaks were ratioed to the isotope corrected internal standard included in each scan type. The inclusion of deuterated and odd chain fatty acids within the internal standard lipids sufficiently mass shifted internal standards away from any biological lipids, therefore allowing accurate and reliable peak intensity measurements to be discerned. Data tables were extracted from Lipidview and imported to Microsoft Excel for cell count normalization, internal standard concentration factoring, statistical analysis, and graphing.

#### 8.4.3 Statistics and error analysis

The mean with 95% confidence intervals was used for error analysis on column-charts throughout and was calculated using Microsoft Excel and conventional equations. For Figure 1, the mean and variance were established using Microsoft Excel to calculate a paired t test. Subsequent t-distribution values were translated to statistical significance (p-values) via relevant degrees of freedom and a critical t-value table. The Fig. 4 heatmap was calculated using R x64 3.6.1 packages and built-in functions (*i.e*., PerformanceAnalytics, Hmisc() and corrplot()). The images of Fig. 5 refer to precursor ions from the same tissue section, except for the *sn-* isomer FDI and the H&E which were conducted on two separate serial sections. Equivalent analyses were conducted on a replicate resected-tumour and an sPLA2-IIa inhibited (KH064) resected tumour, both of which can be found in the supplementary information. Specific statistical details for each of the figures (including number of replicates, p value representation and error model), can be found within the respective figure captions.

## Supplementary information

**Fig. S1:**
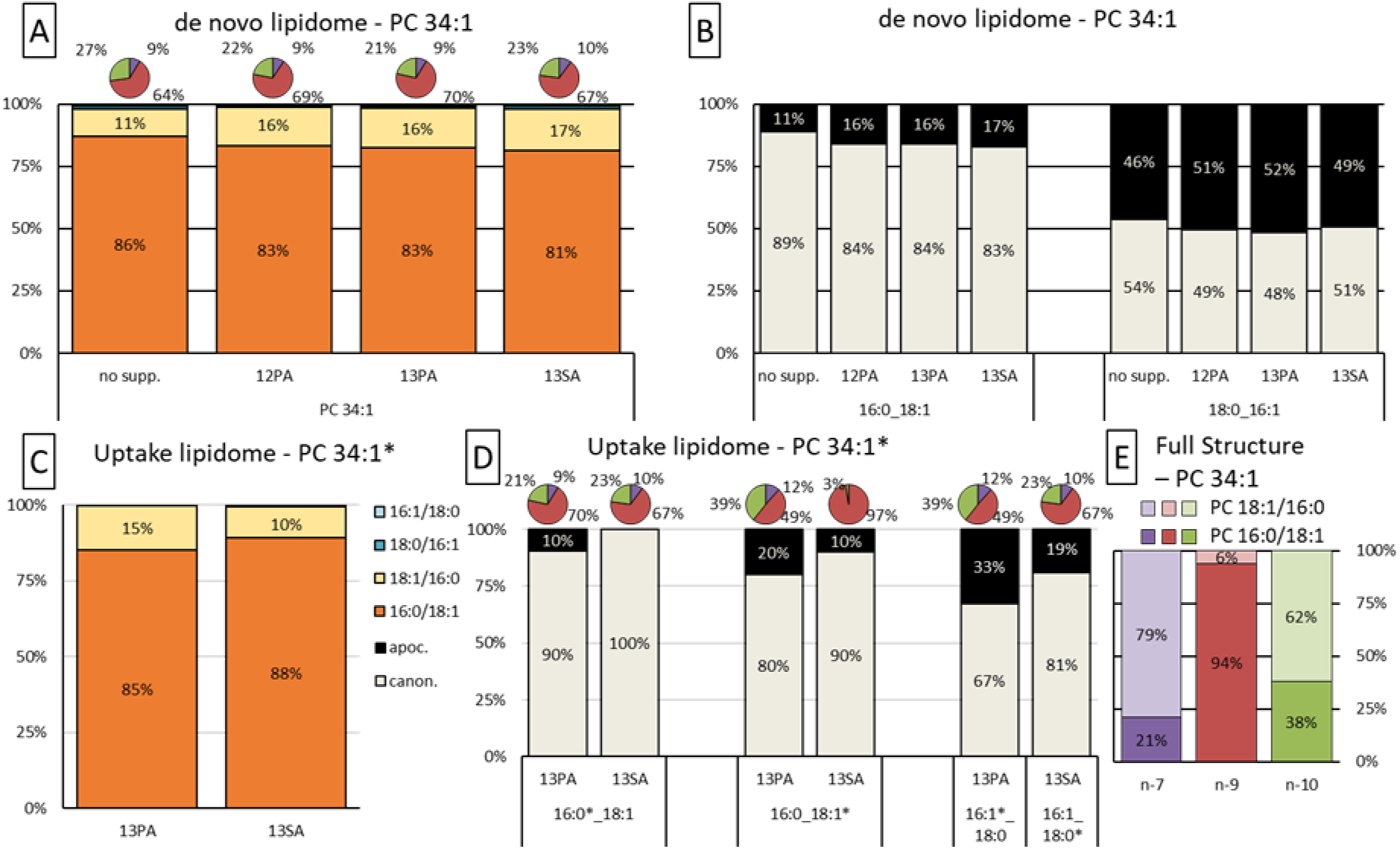
(related to Fig. 2 & 3) *de novo* and *uptake* PC 34:1 DB and *sn-*isomer distributions for non-supplemented LNCaP, 12PA supplemented LNCaP, 13PA supplemented LNCaP and 13SA supplemented LNCaP. **(A)** Relative abundance of FA compositions for *de novo* (unlabelled) PC 34:1, including *sn*-isomer distribution and DB locational isomer distribution (pie charts). **(B)** Relative abundance of *sn*-isomers for either *de novo* (unlabelled) FA composition of PC 34:1. **(C)** Relative abundance of FA compositions for *uptake* (labelled) PC 34:1, including *sn*-isomer distribution. **(D)** Relative abundance of *sn*-isomers for either *uptake* (labelled) FA composition of PC 34:1, including DB locational isomer distribution (pie charts). **(E)** Full structural elucidation of non-supplemented LNCaP PC 34:1, showing combined *sn*- and DB isomer distribution and the association between either type of isomer. Mean ± SEM (95% confidence interval), n = 2, except for all non-supplemented LNCaP data where n = 3.

**Fig. S2:**
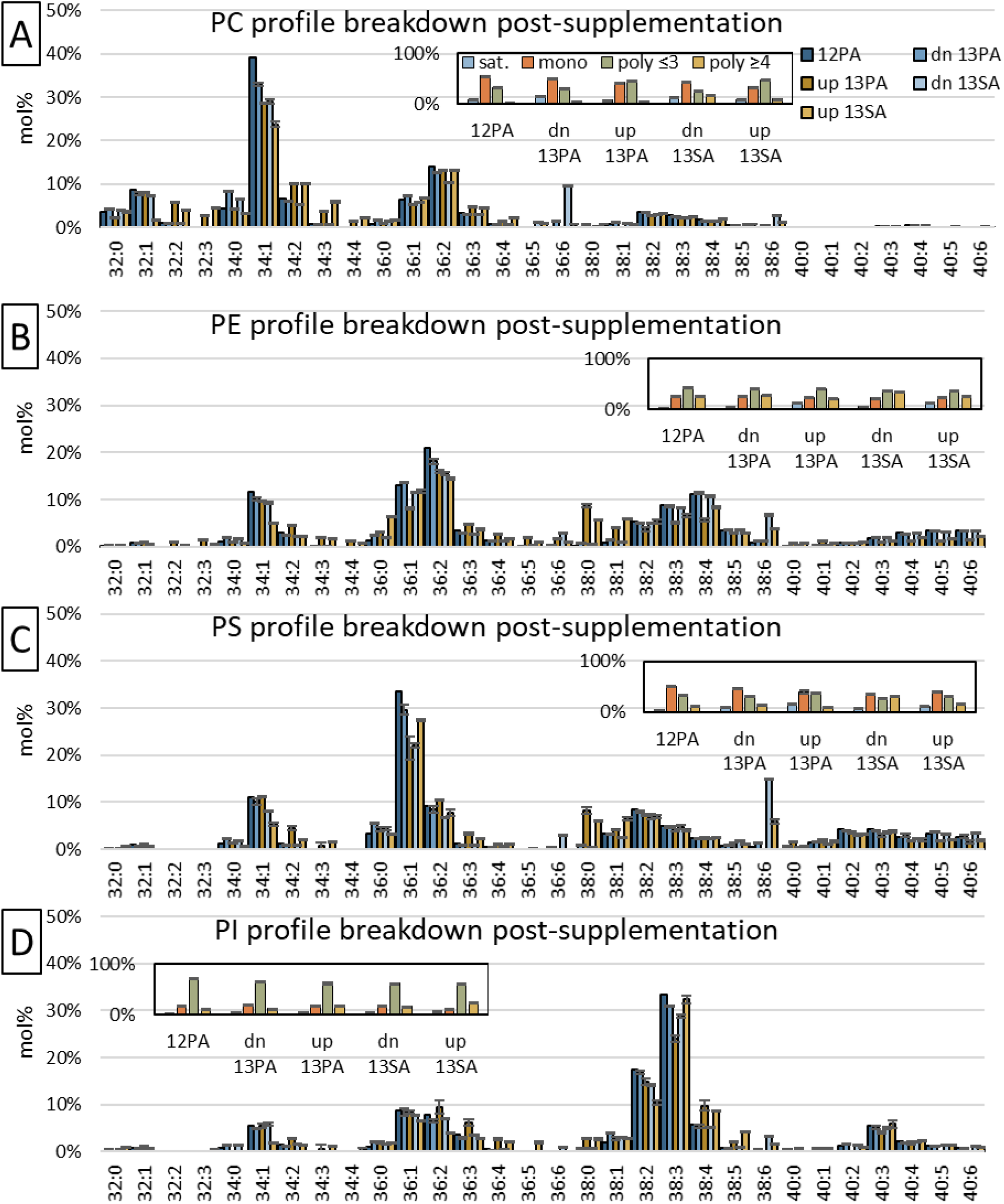
(related to Fig. 1 & 4) Distribution of labelled fatty acids into GPL classes and their effect on species abundance and degree of unsaturation. (A-D) Main bar charts show the species abundance (mol%) for PC, PE, PS and PI and the impact labelled fatty acid supplements indirectly have on the unlabelled lipidome and directly have on the labelled lipidome. Smaller bar charts show the impact supplements have on the degree of saturation for each lipid class. Degree of unsaturation was divided into lipid species that contained fatty acids that were: saturated, monounsaturated, polyunsaturated (≤ 3 DBs) and polyunsaturated (≥4 DBs). Legends are located in panel A; n=2, mean ± SEM (95% confidence interval) displayed.

**Fig. S3:**
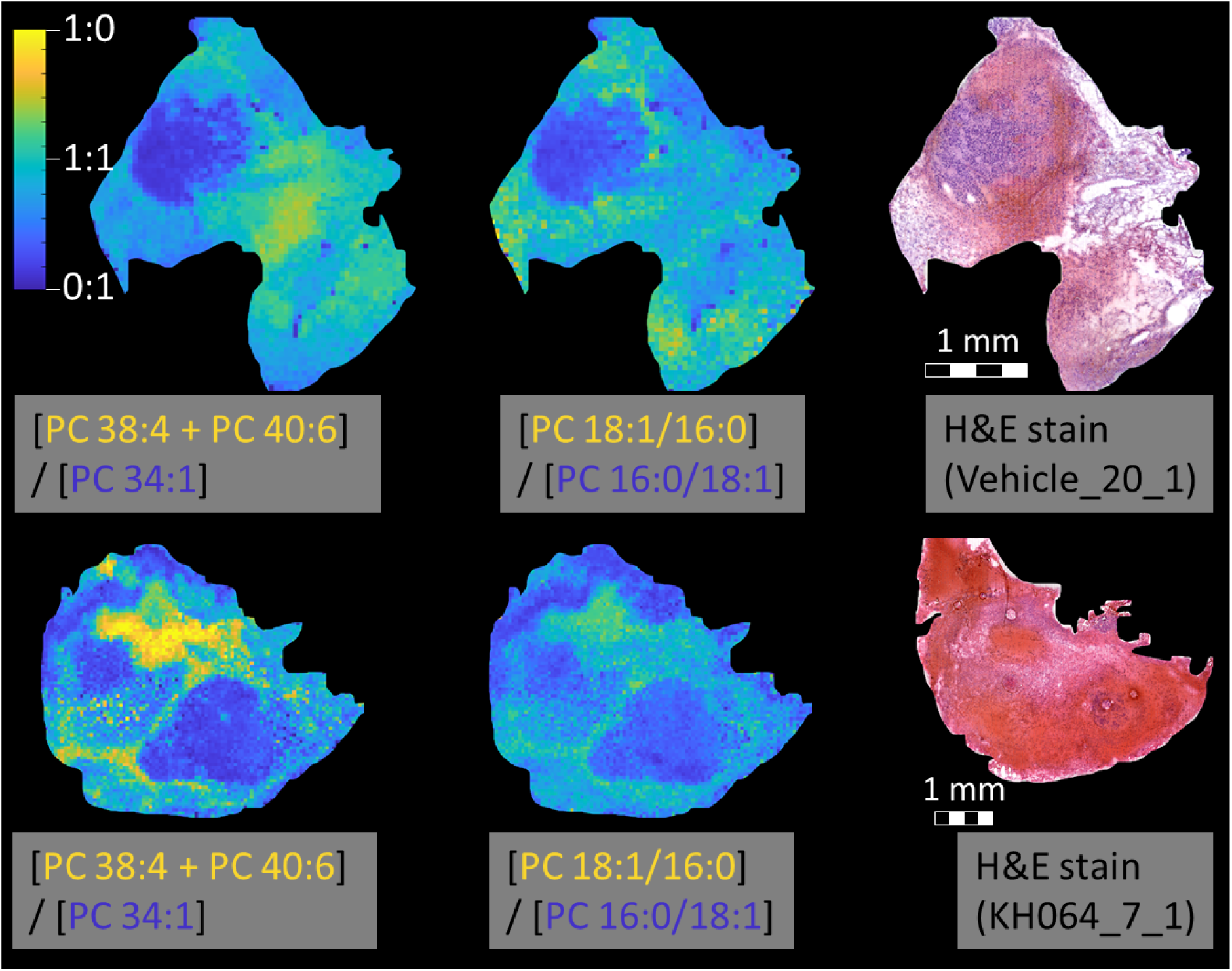
(related to Fig. 5) Uninhibited LNCaP xenograft tissue (biological replicate) and sPLA2-IIA inhibited (KH064) LNCaP xenograft tissue. **(Top row)** Uninhibited (vehicle/treatment-naïve) LNCaP xenograft tissue showing the fractional distribution images (FDI) for PUFA containing PCs and canonomer/apocromer distribution across the tissue. As per the main text figure, positive correlation can be observed between the PUFA species, the PC 18:1/16:0 apocromer and the fatty tissue. **(Bottom row)** As per top row for the KH064 treated xenograft model. An increased abundance of PUFA lipid and decreased abundance of apocromer can be observed across the tissue. Although spatial correlation is still apparent, these changes to lipid species abundance thereby reduces the correlation between either species. Both still appear to be positively correlated with host adipocyte cells. Due to a sampling malfunction, directly adjacent tissue sections could not be analysed, instead the H&E displayed is 3-4 section depths (∼30-40 µm) distal to the MALDI-MSI images.

**Fig. S4:**
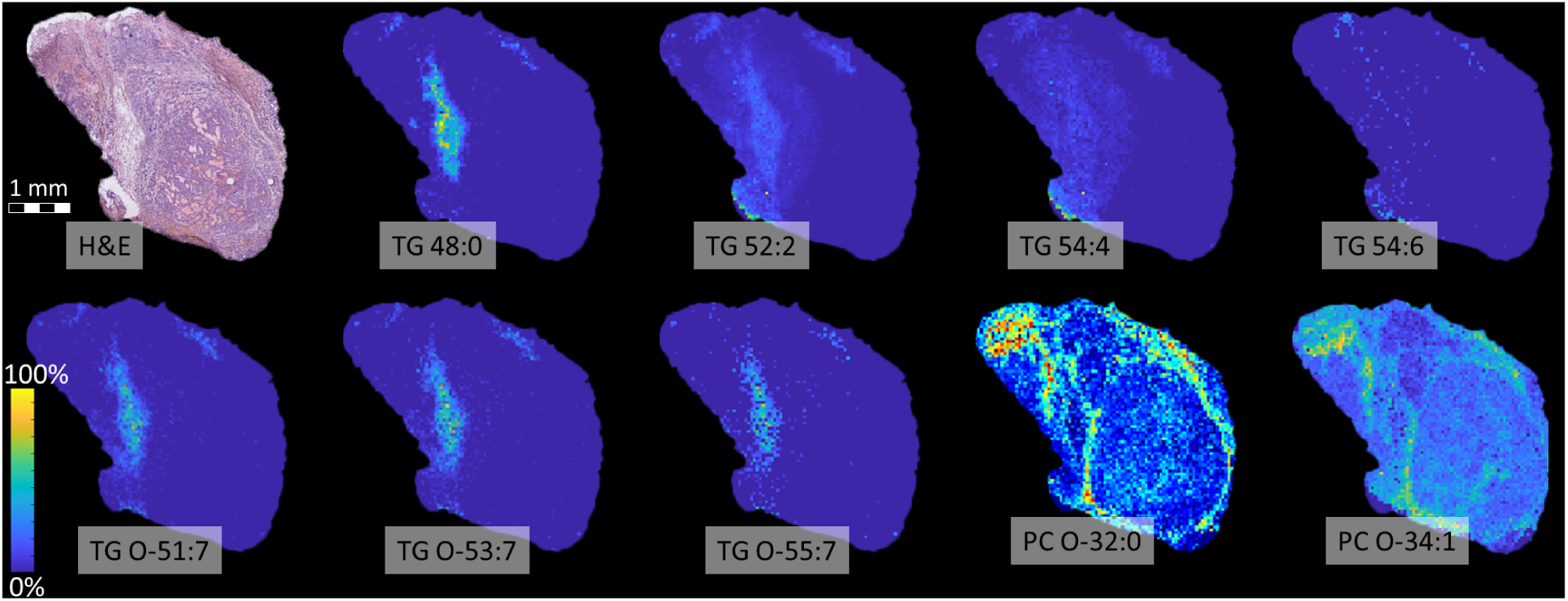
(related to Fig. 5) Ether lipid species present in the uninhibited xenograft tissue. Top row: Distribution and abundance differences of triacylglycerol lipids. Bottom row: Triacylglycerol ethers (TG O-) and phosphatidylcholine ethers (PC O-) were also observed to be spatially correlated to host adipocyte cells, PUFA lipids and increased apocromer abundance (H&E stain). While TG O-appears mostly homogenous across the region, PC O-appears to be more abundant around the perimeter. All species were identified by exact mass (Δppm ≤ 3), with no reasonable alternative identifications found using LIPID MAPS® or other lipid databases. Phosphatidylethanolamine ethers (PE O-) were also potentially identified but were not included due to their isobaric overlap with [PC +K]^+^ precursor ions. All high mass-resolution MALD-MSI images are normalised to self, displaying zero abundance (dark blue) ranging through to highest abundance (orange/red).

